# Distinct role of nucleus accumbens D2-MSN projections to ventral pallidum in different phases of motivated behavior

**DOI:** 10.1101/2020.11.27.401042

**Authors:** Carina Soares-Cunha, Raquel Correia, Ana Verónica Domingues, Bárbara Coimbra, Nivaldo AP de Vasconcelos, Luísa Pinto, Nuno Sousa, Ana João Rodrigues

**Affiliations:** Life and Health Sciences Research Institute (ICVS), School of Medicine, University of Minho, Braga, Portugal; ICVS/3B’s-PT Government Associate Laboratory, Braga/Guimarães, Portugal; Physics Department, Federal University of Pernambuco (UFPE), Recife, Pernambuco, 50670-901, Brazil; Department of Biomedical Engineering, Federal University of Pernambuco (UFPE), Recife, Pernambuco, 50670-901, Brazil

**Keywords:** Nucleus accumbens, ventral pallidum, reward, motivation, medium spiny neurons, D2-MSNs

## Abstract

The nucleus accumbens (NAc) is a key region in motivated behaviors. NAc medium spiny neurons (MSNs) are divided into those expressing dopamine receptor D1 or D2. Classically, D1- and D2-MSNs have been described as having opposing roles in reinforcement but recent evidence suggests a more complex role for D2-MSNs.

Here we show that optogenetic modulation of D2-MSN to ventral pallidum (VP) projections during different stages of motivated behavior has contrasting effects in motivation. Activation of D2-MSN-VP projections during a reward-predicting cue results in increased motivational drive, whereas activation at reward delivery results in decreased motivation; optical inhibition has the opposite behavioral effect. In addition, in a free choice instrumental task, animals prefer the lever that originates one pellet in opposition to pellet plus D2-MSN-VP optogenetic activation, and vice versa for optogenetic inhibition.

In summary, D2-MSN-VP projections play different (and even opposing) roles in distinct phases of motivated behavior.

## Introduction

Organisms are faced with the necessity to look for rewards in the environment to survive. While natural rewards (e.g. food) motivate goal-directed behavior, similar mechanisms appear to be involved in drug seeking, which can lead to drug abuse (Robbins 2002). Interestingly, individuals associate environmental cues with specific rewards, causing for these cues to acquire motivational significance that can shape goal-directed behavior (Rescorla 1994). For example, food-associated cues can induce feeding in sated animals (Holland and Petrovich 2005), and drug-associated cues can elicit drug taking after withdrawal (Grimm et al. 2002), suggesting that such motivational cues have the ability to shape goal-directed behavior. Given these common cue-reward associative mechanisms that natural and non-natural rewards share, by understanding the mechanisms that guide motivated behavior for natural rewards it may be possible to provide insight into similar pathological processes in drug addiction.

The nucleus accumbens (NAc) plays a significant role in regulating reward-seeking and motivated behaviors (Berridge 2007). Electrophysiological studies have shown that NAc neurons encode both the predictive value of environmental stimuli and the specific motor behaviors required to respond to them (Carelli 2002; Nicola et al. 2004). Interestingly, with learning, NAc neurons, and in particular those located in the core subregion (Ambroggi et al. 2011), develop responses to cues predicting rewards (Roitman et al. 2005). In addition, others have shown that the NAc contains distinct classes of neurons: one that increases firing at cue exposure in a cue-reward association task; while other exhibits attenuated firing rate specifically at reward delivery (Gale et al. 2014).

The NAc receives dopamine signals from the ventral tegmental area (VTA), which acts predominantly via activation of D1 or D2 dopamine receptors that are expressed by largely non-overlapping populations of medium spiny neurons (MSNs) (Gerfen and Surmeier 2011). These two MSN sub-populations project to different outputs: D1-MSNs project to the VTA, while both D1- and D2-MSNs project to the ventral pallidum (VP) (Lu et al. 1998; Kupchik et al. 2015). Initial studies suggested that different NAc MSN subtypes play distinct and opposing roles in motivated behaviors: while activation of D1-MSNs promotes reward-related outcomes, activation of D2-MSNs blunts rewarding events (Lobo et al. 2010; Kravitz et al. 2012; Calipari et al. 2016). However, other studies challenged this functional opposing view and showed that NAc D2-MSN optical stimulation promotes self-stimulation (Cole et al. 2018), suggesting that D2-MSNs may be pro-rewarding in some contexts. Moreover, we have shown that brief optical activation of D2-MSNs paired with a reward-predicting cue enhances motivation to obtain food rewards (Soares-Cunha et al. 2016, 2018). In addition, specific cue-driven activation of D2-MSN-VP projections would increase motivation towards obtaining a food reward (Soares-Cunha et al. 2018). Interestingly, this behavioral effect was triggered by a transient decrease in activity of GABAergic neurons within the VP which, in turn, resulted in disinhibition of dopaminergic activity of the VTA (Soares-Cunha et al. 2018), known to induce increased motivational levels (Ilango et al. 2014; Han et al. 2017). Nonetheless, others have shown that chronic inhibition of D2-MSNs during a progressive ratio (PR) task (using chemogenetics), leads to enhanced motivation without affecting the sensitivity to reward devaluation (Carvalho Poyraz et al. 2016). Interestingly, despite increasing motivation, inhibition of D2-MSNs would cause animals to initiate more frequently behavior without goal-directed efficiency (Gallo et al. 2018), an effect caused by disinhibition of VP GABAergic activity (Gallo et al. 2018). These studies and others further support the notion that reward-related behavioral responses depend on normal connectivity between NAc and VP (Chang et al. 2018), and that VP is a critical interface between reward processing and motor output (Chang et al. 2018; Ottenheimer et al. 2018).

In general, these results seem to point for the importance of D2-MSN-VP in specific sequences of behavior such as action initiation and cue-driven responses. Thus, to better understand the contribution of D2-MSN-VP projections in reward-related behaviors, we optically manipulated these neurons during specific segments of behavior. We show that optogenetic stimulation of D2-MSN-VP projections can bi-directionally modulate motivated behavior depending on the timing of stimulation: activation during a cue that predicts a reward increases motivation while activation of the same neurons at reward delivery decreases motivation in different behavioral paradigms. This study adds to a better understanding on the role of D2-MSN-VP projections in motivated behaviors and shows that these neurons play a much more complex role in reinforced behavior than previously anticipated.

## Results

### Optogenetic activation D2-MSN terminals modulates ventral pallidum activity

Previous data from our group showed that optogenetic stimulation of NAc core D2-MSNs cell bodies during cue exposure increases motivation and is rewarding (Soares-Cunha et al. 2016). However, it remains to be determined if this behavioral effect is through D2-MSN terminals in the VP or through another downstream region. So, in order to better understand the role of D2-MSN-VP projections in motivated behavior, we used optogenetics to manipulate these terminals.

We unilaterally injected an AAV5 containing a construct with channelrhodopsin (ChR2) in fusion with enhanced yellow fluorescent protein (eYFP) under the control of the dopamine receptor D2 minimal promoter (pAAV-D2R-hChR2(H134R)-eYFP) in the NAc core of wild type *Wistar Han* rats (D2-ChR2 group; Figure 1A), which allows specific manipulation of D2^+^ neurons (Soares-Cunha et al. 2016; Zalocusky et al. 2016). Expression of eYFP was observed in cell bodies in the NAc core (Figure 1B) and in D2-MSN terminals located in the dorsal VP (dVP; Figure 1C), the VP sub-region that is innervated by the NAc core D2-MSNs (Heimer et al. 1991). We next used single-cell *in vivo* electrophysiology in anesthetized animals to evaluate optically-evoked response of dVP neurons to D2-MSN terminal activation (1s, 25ms light pulses at 20Hz) (Figure 1D). Half of recorded dVP neurons responded to D2-MSN terminal stimulation by decreasing firing rate (15 cells), 10% increased firing rate (3 cells) and 40% (12 cells) presented no change in comparison with baseline (Figure 1E). As a consequence, the net firing rate of dVP significantly decreased during the stimulus period (*F*_(2,29)_ = 7.32, *p* = 0.0083, *one way* ANOVA; Figure 1E) in comparison to baseline activity (Bonferroni *post hoc, p* = 0.0339). After optical stimulation, dVP firing rate returned to baseline levels (Bonferroni *post hoc, p* = 0.6083). This effect is also shown in the temporal variation of activity of all recorded neurons (Figure 1F).

**Figure 1.**
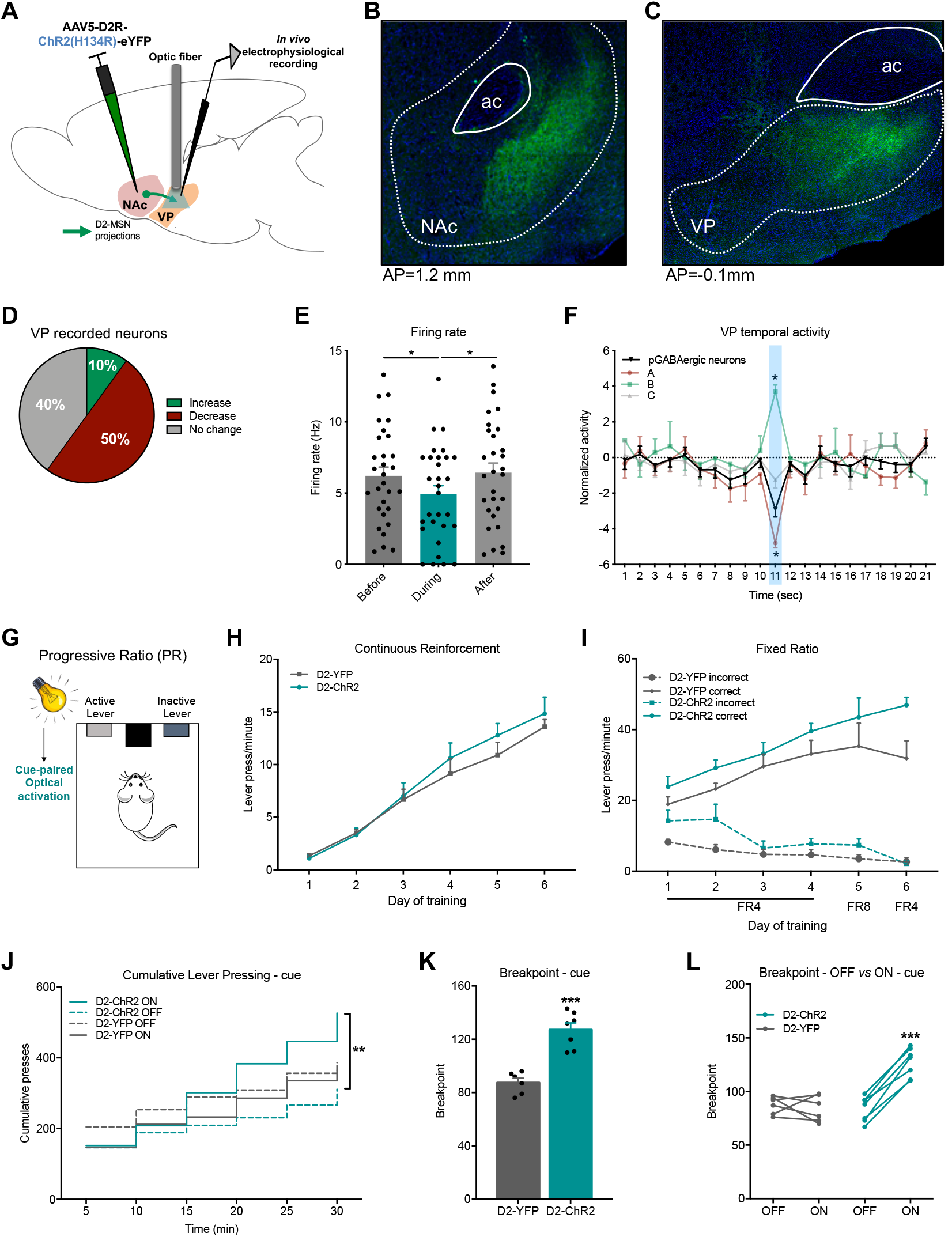
with 1 supplement. Optogenetic activation of D2-MSN-VP terminals during cue exposure increases motivation. ***A*** Strategy used for NAc D2-MSN-VP projection optogenetic stimulation and electrophysiological recordings in the dVP. ChR2-YFP was unilaterally injected into NAc, mainly targeting core. Stimulation was made in terminals in the dVP. ***B*** Representative immunofluorescence showing eYFP expression in the NAc and ***C*** in terminals in the dVP; scale bar=400μm; AP = anteroposterior. ***D*** Pie chart showing that 50% of putative GABAergic (pGABAeric) VP neurons decrease firing rate during optical stimulation (20 pulses of 25ms at 20Hz). ***E*** dVP neurons significantly decrease firing rate in response to optical stimulation of D2-MSNs terminals (n=30 neurons/4 rats). ***F*** PSTH of temporal variation of the normalized activity of pGABAeric VP neurons that decrease (A; green, n=15 neurons), increase (B; red, n=3 neurons) and do not change activity (C; gray, n=12 neurons) during the stimulation period (blue). ***G*** Rats were submitted to a PR session in which optogenetic activation of D2-MSN-VP (20 pulses of 25ms at 20Hz) was paired with cue light presentation above the active lever at trial initiation. ***H*** CRF training sessions of the PR schedule, shown as average of number of lever presses per minute. ***I*** FR training sessions of the PR test shown as average number of lever presses per minute. ***J*** Optogenetic stimulation of D2-MSN-VP projections during cue exposure increases cumulative presses in the PR test session. ***K*** D2-MSN-VP terminal stimulation induces a significantly higher breakpoint in comparison to D2-YFP animals. ***L*** All D2-ChR2 rats increase breakpoint in the session with optical stimulation (ON) in comparison with the session without (OFF). n_D2-ChR2_=7, n_D2-eYFP_=6. Error bars denote SEM. \**p* ≤ 0.05.

### Cue-paired optogenetic activation of D2-MSN-VP projections increases motivation

Previous data presented conflicting results regarding D2-MSNs’ role in motivation (Carvalho Poyraz et al. 2016; Soares-Cunha et al. 2016, 2018; Gallo et al. 2018), suggesting that these neurons play distinct roles in different stages of motivated behavior. In this work, we decided to focus on the modulation of D2-MSN-VP terminals in motivated behaviors. First, we tested D2-ChR2 animals in the PR task that has one active lever associated with reward delivery and one inactive lever (Figure 1G). Each trial begins with turning ON of a cue light located above the active lever. The PR directly measures motivational level which is given by the breakpoint, that is the moment when the animal gives up pressing for the reward (Wanat et al. 2010). Control group was injected with an AAV containing pAAV-D2R-eYFP (D2-YFP) (Figure 1 – figure supplement 1A-B).

During continuous reinforcement (CRF) training, both groups increased lever pressing throughout days similarly (*F*_(1,66)_ = 1.5, *p* = 0.2230, *two way* ANOVA; Figure 1H). All animals increased lever pressing in the fixed-ratio (FR) schedule days in the correct *versus* incorrect lever (*F*_(3,22)_ = 61.3, *p* < 0.0001, *two way* ANOVA; Figure 1I).

After stable lever pressing, we performed optical stimulation (1s, 25ms pulses at 20Hz) of D2-MSN terminals in the dVP paired with cue light period of the PR session. D2-MSN-VP stimulation induced a significant increase in the number of cumulative lever presses (D2-ChR2 ON *versus* D2-ChR2 OFF; *F*_(5,72)_ = 10.6, *p* < 0.0001, *two way* ANOVA; Figure 1J). D2-ChR2-stimulated animals presented a 51% increase in breakpoint in comparison to D2-YFP-stimulated rats (*t*_11_= 5.98, *p* < 0.0001, unpaired *t* test; Figure 1K). All D2-ChR2 rats displayed a significant increase in breakpoint in the session with optical stimulation (ON) in comparison with the session without optical stimulation (OFF) (*t*_6_= 10.2, *p* < 0.0001, paired *t* test; Figure 1L), which was counterbalanced between days. This enhancement was not due to changes in food consumption, since the number of pellets earned was similar in the two PR sessions (*t*_6_= 2.0, *p* = 0.0930, paired *t* test; Figure 1 – figure supplement 1C).

When optogenetic activation occurred during the inter-trial interval (ITI), no differences in the cumulative lever pressing (Figure – figure supplement 1D), breakpoint (Figure – figure supplement 1E), or number of food pellets earned (Figure 1 – figure supplement 1F) was observed between D2-ChR2 and D2-YFP animals, showing that the motivation boost effect is restricted to particular stages of behavior.

### Cue-paired optogenetic inhibition of D2-MSN-VP projections decreases motivation

We next inhibited D2-MSN-VP terminals during PR performance. Animals were injected with an AAV5 containing halorhodopsin (eNpHR) under the control of the D2 minimal promoter (pAAV-D2R-eNpHR3.0-eYFP) (Figure 2A,B; Figure 1 – figure supplement 1A,B) (Soares-Cunha et al. 2016).

**Figure 2.**
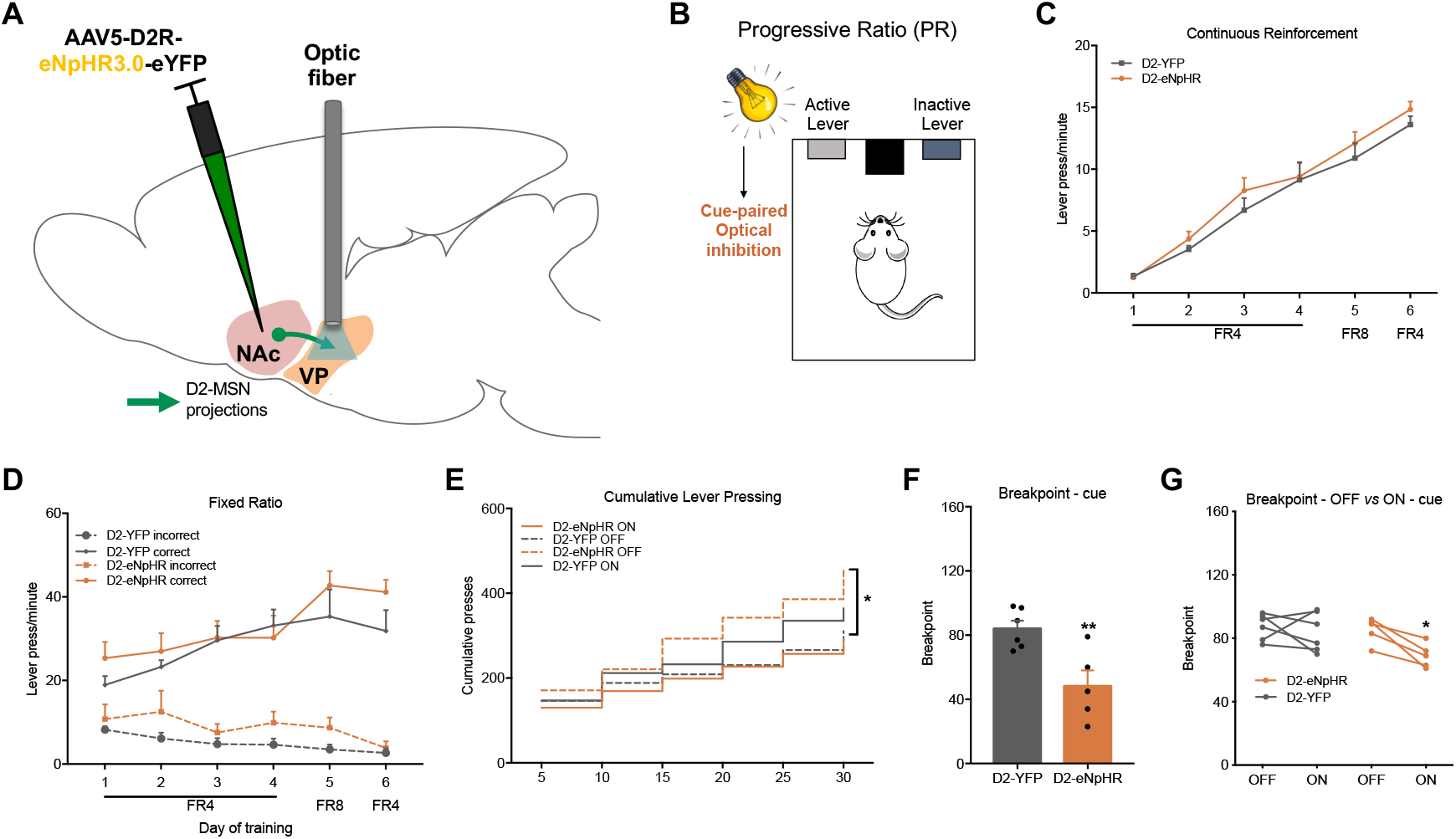
with 1 supplement. Optogenetic inhibition of D2-MSN-VP terminals during cue exposure decreases motivation. ***A*** Strategy used for optogenetic inhibition of D2-MSN-VP terminals. ***B*** Rats were submitted to a PR session in which optogenetic inhibition of D2-MSN-VP (10sec of constant light at 10 mW) was paired with cue light presentation above the active lever at trial initiation. ***C*** CRF training sessions of the PR schedule, shown as average of number of lever presses per minute. ***D*** FR training sessions of the PR test shown as average number of lever presses per minute. ***E*** Optogenetic inhibition of D2-MSN-VP terminals during cue exposure decreases cumulative presses in the PR test session. ***F*** D2-MSN-VP terminal inhibition induces a significantly lower breakpoint in comparison to D2-YFP animals. ***G*** All D2-ChR2 rats decrease breakpoint in the session with optical inhibition (ON) in comparison with a session without optical inhibition (OFF). n_D2-eNpHR_=5, n_D2-eYFP_=6. Error bars denote SEM. \**p* ≤ 0.05; \*\**p* ≤ 0.01.

D2-NpHR and D2-YFP rats presented similar rate of lever pressing in the training sessions. During CRF training, both groups increased lever pressing similarly across days of training (*F*_(1,60)_ = 2.9, *p* = 0.0962, *two way* ANOVA; Figure 2C), and all animals increased lever pressing in the FR schedule in the correct *versus* incorrect lever (*F*_(3,18)_ = 27.5, *p* < 0.0001, *two way* ANOVA; Figure 2D). D2-MSN-VP optical inhibition (10s constant light at 10mW) during cue light period of the PR session, induced a significant decrease in the number of cumulative lever presses (D2-eNpHR ON *versus* D2-eNpHR OFF; *F*_(5,48)_ = 5.3, *p* = 0.0006, *two way* ANOVA; Figure 2E). This was translated into a 43% decrease of the breakpoint of D2-eNpHR group in comparison to D2-YFP stimulated rats (*t*_9_=3.4, *p* = 0.0076, unpaired *t* test; Figure 2F). All D2-NpHR rats displayed a significant decrease in breakpoint in the ON session in comparison with the OFF session (*t*_6_=10.2, *p* < 0.0001, paired *t* test; Figure 2G). D2-NpHR and D2-YFP rats earned a similar number of pellets in the PR session (*t*_9_=0.1, *p* = 0.8925, unpaired *t* test; Figure – figure supplement 1A).

Optogenetic inhibition during the ITI did not induce differences in the cumulative lever pressing (Figure 2 – figure supplement 1B), breakpoint (Figure 2 – figure supplement 1C), or number of food pellets earned (Figure 2 – figure supplement 1D).

### Impact of reward-paired optogenetic modulation of D2-MSN-VP projections in motivation

D2-MSN-VP modulation during cue exposure induced robust changes in motivational drive, but the impact of this manipulation during other task-relevant periods remained unknown. Thus, we next paired D2-MSN-VP projection activation/inhibition with cue exposure as in the previous task, and in another session, optical modulation was paired with reward (pellet) delivery (Figure 3A). Interestingly, D2-ChR2 animals presented a significantly lower number of cumulative lever presses (30.2% less) performed in the PR session with optical stimulation paired with reward delivery, in comparison with D2-YFP animals (*F*_(5,60)_ = 9.9, *p* < 0.0001, *two way* ANOVA; D2-ChR2 reward *versus* D2-YFP reward, *Tukey’s* multiple comparison, *p* < 0.0001; Figure 3B). In addition, D2-ChR2 animals presented a significantly lower number of cumulative lever presses in comparison to the PR session in which they received optical activation at cue exposure (48.2% decrease; *Tukey’s* multiple comparison, *p* < 0.0001; Figure 3B). As compared to D2-YFP, there was a 46.5% reduction in breakpoint (*t*_19_=7.1, *p* < 0.0001, unpaired *t* test); and a 64% reduction in comparison to D2-ChR2 cue session (*t*_11_=15.6, *p* < 0.0001, paired *t* test; Figure 3C,D).

**Figure 3.**
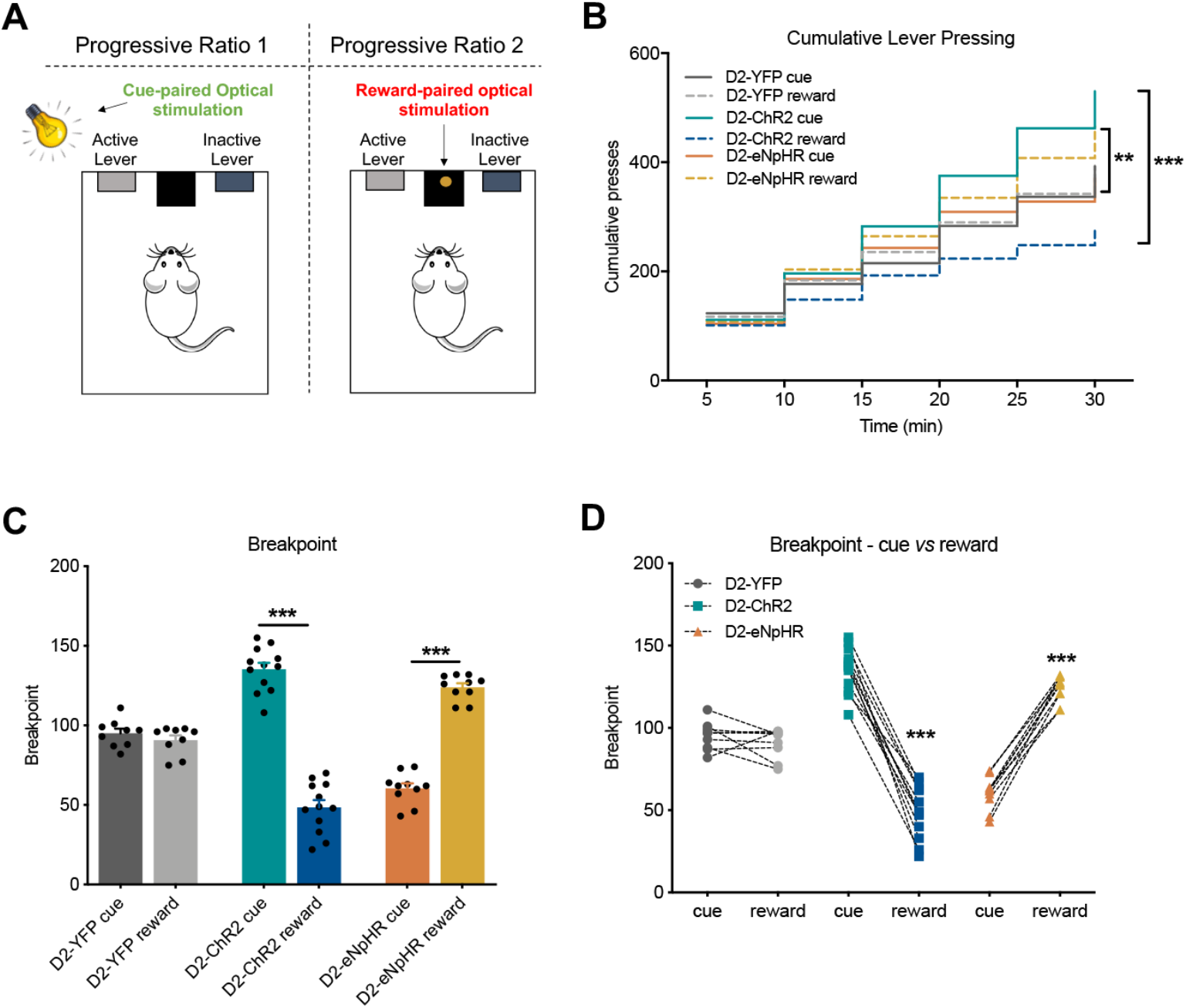
with 1 supplement. Optogenetic modulation of D2-MSN-VP terminals at reward delivery decreases motivation. ***A*** Rats were tested in two PR sessions: in one, optogenetic modulation – excitation: 20 pulses of 25ms at 20Hz; inhibition: 10sec of constant light at 10 mW - was paired with cue presentation; in the other session, optogenetic modulation was paired with reward delivery. ***B*** Optogenetic activation of D2-MSN-VP terminals at cue exposure results in a significantly higher number of cumulative lever presses in comparison to D2-YFP control animals and in comparison with D2-ChR2 stimulated during reward delivery; optogenetic inhibition of D2-MSN-VP terminals at cue exposure results in a significantly lower number of cumulative lever presses in comparison to D2-YFP animals and in comparison with D2-eNpHR stimulated during reward delivery. ***C, D*** Optogenetic stimulation of D2-MSN-VP terminals at cue exposure results in a significantly higher breakpoint in comparison with control animals and with optogenetic stimulation of the same neurons at reward delivery; the opposite is observed with optogenetic inhibition of the same projections. n_D2-ChR2_=12; n_D2-eNpHR_=11, n_D2-eYFP_=10. Error bars denote SEM. \*\**p* ≤ 0.01; \*\*\**p* ≤ 0.001.

Optogenetic inhibition of D2-MSN-VP projections paired with reward delivery significantly increased the number of cumulative lever presses in comparison to both D2-YFP (18.2% increase; *Tukey’s* multiple comparison, *p* = 0.0495; Figure 3B) and D2-eNpHR cue (24.3% increase; *Tukey’s* multiple comparison, *p* = 0.0036; Figure 3B). As compared to D2-YFP, there was a 36.9% increase in breakpoint (*t*_17_=8.6, *p* < 0.0001, unpaired *t* test); and a 105% increase in comparison to D2-eNpHR cue (*t*_10_=11.9, *p* < 0.0001, paired *t* test; Figure 3C,D).

No significant differences were observed in the number of food pellets earned during the PR sessions (Figure 3 – figure supplement 1A).

### Optogenetic activation of D2-MSN-VP projections paired with reward shifts preference and decreases motivation

Considering the distinct role of D2-MSN-VP projections in different stages of the PR test, we tested animals in another reward-related task. Animals were tested in a two-lever free choice behavioral paradigm, in which animals have available two levers that deliver one food pellet each; one lever is randomly assigned to stimulate D2-MSN-VP projections (stim+), while the other lever is assigned to deliver the pellet alone (stim-) (Figure 4A).

**Figure 4.**
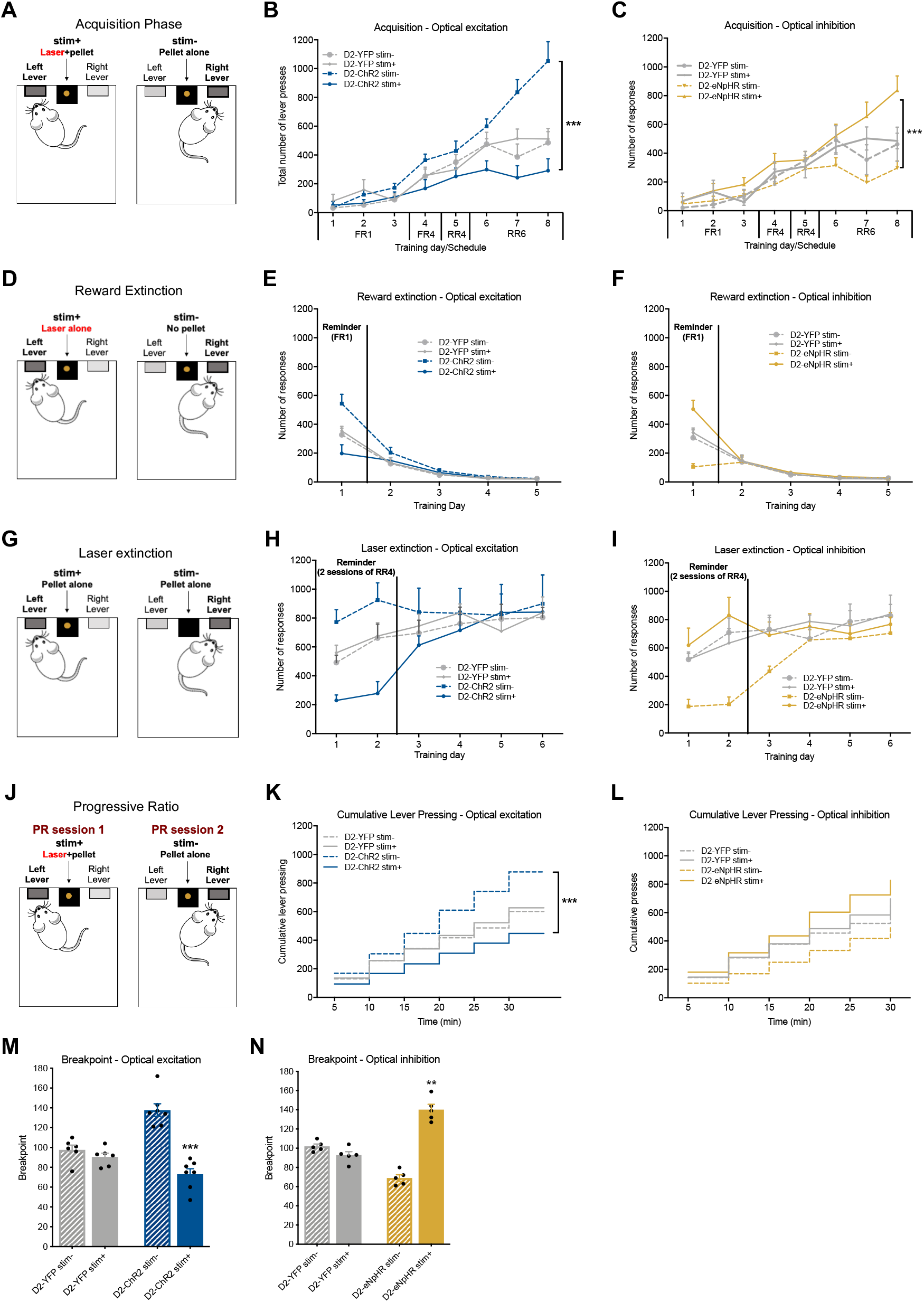
with 2 supplement2. Optical activation/inhibition of D2-MSN-VP terminals paired with reward delivery reduces/increases preference and decreases motivation. ***A*** In a two-choice acquisition lever pressing task, pressing stim+ lever results in the delivery of a food pellet reward and optical stimulation (D2-ChR2; 20 pulses of 25ms at 20Hz) or optical inhibition (D2-eNpHR; 10sec of constant light at 10 mW); pressing stim- lever results in delivery of food pellet reward alone. ***B*** Time-course representation of the responses of D2-ChR2 and D2-eYFP rats. D2-ChR2 rats show reduced preference for stim+ lever in comparison with stim- lever, while D2-eYFP show no preference for either lever. ***C*** Time-course representation of the responses of D2-eNpHR and D2-eYFP rats; D2-eNpHR rats show increased preference for stim+ lever in comparison with stim- lever, while D2-eYFP show no preference. ***D*** Next, rats were tested in pellet extinction sessions, in which no reward is given in any of the levers. ***E*** In pellet extinction, all D2-ChR2 decrease response for both levers. ***F*** In pellet extinction, all D2-eNpHR animals decrease response for both levers. ***G*** In laser extinction sessions, pressing in either stim+ or the stim- results in the delivery of food pellet alone, making the reward in both levers equal. ***H*** In laser extinction conditions, D2-ChR2 rats showed no preference for either lever; the same was observed for D2-eYFP rats. ***I*** under laser extinction, D2-eNpHR rats showed no preference for either lever, pressing the same amount of times in both; the same was observed for D2-eYFP rats. ***J*** Rats were subjected to two PR sessions, one for each lever: in one session, animals are tested for the stim+ lever, and in the other session animals are tested for the stim- lever. ***K*** Cumulative lever pressing, showing that D2-ChR2 rats press less on the stim+ lever session in comparison with the stim- lever session. ***L*** Cumulative lever pressing, showing that D2-eNPHR rats press more on the stim+ lever session in comparison with the stim- lever session. ***M*** Decrease in the breakpoint for stim+ session in comparison with stim- session in D2-ChR2 animals. ***N*** Increase in breakpoint for stim+ session in comparison with stim- session for D2-eNpHR rats. n_D2-ChR2_=7, n_D2-eYFP_=6. Error bars denote SEM. \*\*\**p* ≤ 0.001.

Throughout acquisition days, D2-ChR2 rats showed a clear preference for pressing the stim- lever in comparison with the stim+ lever (*F*_(7,48)_ = 7.3, *p* < 0.0001, *two way* ANOVA; *Bonferroni* multiple comparison, *p* < 0.0001; Figure. 4B). As expected, D2-YFP rats showed no preference for either lever (Figure 4B). Regardless of this preference, no significant differences were found in the total number of lever presses (in both levers) in the last day of training between D2-ChR2 (average of 1230.1 lever presses) and D2-YFP (average of 996.3 lever presses) rats (*t*_11_ = 1.6, *p* = 0.1426, unpaired *t* test; data not shown).

Considering this preference, we next decided to evaluate individuals’ behavior in reward extinction conditions. The same behavioral task was performed but no pellet was delivered on either lever, though optical stimulation still occurred at stim+ lever (Figure 4D). Interestingly, both groups decrease lever pressing in both levers, indicating that the instrumental response was dependent on the delivery of the reward (Figure 4E).

After 2 reminder sessions with laser paired with reward delivery in the assigned lever, we performed the task in laser extinction conditions, i.e., the outcome is the same in both levers – one pellet – but no stimulation is given (Figure 4G). D2-ChR2 animals showed no preference for either lever under laser extinction conditions (*F*_(5,30)_ = 1.2, *p* = 0.3178, *two way* ANOVA; *Bonferroni* multiple comparison, p > 0.9999; Figure 4H).

Next, we performed two sessions of PR, one session for each lever (Figure 4J). Interestingly, D2-ChR2 rats pressed cumulatively more in the PR session in which animals worked for the stim- lever, in comparison with the session in which they had to press for the stim+ lever (*F*_(6,72)_ = 135.7, *p* < 0.0001, *two way* ANOVA; *Bonferroni* multiple comparison, *p* < 0.0001; Figure 4K). There was a significant decrease in breakpoint (53%) in all D2-ChR2 animals for stim+ lever in comparison to stim- (*t*_6_ = 10.3, *p* < 0.0001, paired *t* test; Figure 4M). No differences in the number of food pellet consumed during both PR sessions was found (D2-ChR2 stim+ *versus* D2-ChR2 stim-; *t*_6_ = 0.6, *p* = 0.5729, paired *t* test; Figure 4 – figure supplement 1A).

### Optogenetic inhibition of D2-MSN-VP projections paired with reward shifts preference and increases motivation

We next performed the two-lever free choice behavioral paradigm with optogenetic inhibition of D2-MSN-VP terminals. Rats were presented with two levers that originated a food pellet, but stim+ was now associated with optogenetic inhibition of D2-MSN-VP projections. A significant difference was found between choice of levers along the days of training: D2-eNpHR rats showed a significant preference for pressing the stim+ lever in comparison with stim- lever (*F*_(7,56)_ = 26.8, *p* < 0.0001, *two way* ANOVA; Figure 4C). As anticipated, D2-YFP rats showed no preference for either lever (Figure 4C). No significant differences were found in the total number of lever presses in the last day of training between D2-NpHR (average of 1135 lever presses) and D2-YFP (average of 996.3 lever presses) (*t*_9_ = 0.9, *p* = 0.3780, unpaired *t* test; data not shown).

We next performed the task in food extinction conditions (Figure 4D). Both groups significantly decreased lever pressing on either lever (Figure 4F), indicating that stimulation alone is not sufficient to maintain increased lever pressing.

Next, we performed the same task but in laser extinction conditions, which makes the outcome (pellet alone) equal in both levers (Figure 4G). D2-eNpHR animals showed no preference for either lever (*F*_(5,40)_ = 6.8, *p* = 0001, *two way* ANOVA; *Bonferroni* multiple comparison, *p* > 0.9999; Figure 4I).

After performing a reminder of the two-lever free choice task, we performed two PR sessions, one for each lever (Figure 4J). D2-eNpHR rats pressed cumulatively more in the PR session in which animals worked for the stim+ lever in comparison with stim- lever (*F*_(5,40)_ = 217.0, *p* < 0.0001, *two way* ANOVA; *Bonferroni* multiple comparison, *p* < 0.0001; Figure 4L). This was reflected in a significantly increased breakpoint (103%) in all D2-eNpHR animals for the PR session of stim+ lever in comparison with the session of stim- lever (*t*_4_ = 8.6, *p* = 0.0010, paired *t* test; Figure 4N). No differences in the number of food pellets consumed in both PR sessions was observed (D2-eNpHR stim+ versus D2-eNpHR stim-; *t*_4_ = 2.1, *p* = 0.1045, paired *t* test; Figure 4 – figure supplement 1B).

### No differences in food consumption with D2-MSN-VP modulation

In order to further verify if D2-MSN-VP projections play a relevant role in food consumption we evaluated the amount of food – normal chow and food pellets – that animals would consume in one session with no optogenetic modulation and one session with either optogenetic activation (D2-ChR2) or optogenetic inhibition (D2-eNpHR) of D2-MSN-VP projections. For this, all animals performed three days of free food consumption in a cage similar to the home cage and the amount of food consumed was monitored by evaluating the weight at the end of each 30-minute session; all animals performed two days of consumption test without optical stimulation (that were averaged) and one day with optogenetic stimulation. Neither optogenetic activation nor optogenetic inhibition of D2-MSN-VP projections had a significant impact in the consumption of normal chow or food pellets (Figure 4 – figure supplement 2A-F), indicating that the changes in motivation caused by optogenetic modulation were not due to differences in food consumption or satiety.

## Discussion

Despite remarkable advances in identifying the role of specific neuronal populations of the reward circuit in motivated behaviors (Parker et al. 2016; Saunders et al. 2018; Engelhard et al. 2019), there is still a lot of controversy regarding NAc D1- and D2-MSNs (Soares-Cunha et al. 2016, 2019; Natsubori et al. 2017; Cole et al. 2018). D2-MSNs have been considered to play a crucial role in inducing transient punishment and aversive responses (Kravitz et al. 2012); however, accumulating evidence points to a heterogenous role of this neuronal population in behavior (Soares-Cunha et al. 2016, 2019; Natsubori et al. 2017; Cole et al. 2018). In the present study we report that optogenetic activation of D2-MSN-VP projections occurring at the same time as reward-predicting cue increases motivation. Contrary, if optogenetic stimulation was paired with reward (pellet) delivery, this resulted in decreased breakpoint/motivation. Inhibition experiments led to opposit effects in behavior, confirming these results. In addition, we report that in a free-choice task in which rats could press a lever to receive a food pellet or a food pellet paired with laser stimulation, a shift in preference towards pellet alone was observed, and *vice versa* for optogenetic inhibition. These results suggest that D2-MSN terminals in the VP are likely differentially activated during different phases of motivated behaviors.

Previously we showed that cue-paired optogenetic activation of D2-MSNs in the NAc leads to a significant increase in motivation (Soares-Cunha et al. 2016). However, other studies showed that chronic inhibition of these neurons using chemogenetics increases motivation (Carvalho Poyraz et al. 2016; Gallo et al. 2018). This apparently contradictory findings can now be reconciled with the evidence that D2-MSNs appear to play distinct roles during different stages of reward-related tasks. Our data indicates that D2-MSN-VP projections are important to add value to a cue that predicts a future reward, and to increase effort towards obtaining the reward. Conversely, activation of D2-MSN-VP projections at reward delivery decreases breakpoint/motivation, contrary to the observed effects with cue-paired optical activation. This behavioral data suggests that D2-MSN-VP inputs activation is necessary to invigorate cue value, but suppression of D2-MSN-VP activity is required for efficient reward consumption.

Altogether the PR task data pinpoints that D2-MSNs projecting to the VP are necessary to encode the value of a reward-predicting cue, but not so much for the reward itself. In agreement, in the two-choice task, optogenetic activation of D2-MSN-VP projections paired with reward delivery (stim+) shifts preference for stim- lever; the contrary was observed with optical inhibition of these terminals. Interestingly, and in support of our behavioral results, data from extracellular recordings in the dorsomedial striatum has shown that D2-MSNs are selectively active during action initiation, but are suppressed at outcome delivery (Nonomura et al. 2018). In addition, electrophysiological studies performed in the NAc have shown that MSNs exhibit phasic increases in firing rate during cue presentations, but attenuated firing rates at reward delivery (Gale et al. 2014). Similarly, electrophysiological recordings in the NAc during a Pavlovian conditioning approach task showed that MSNs exhibit significant responses during task performance: while MSNs increase firing rate during conditioned stimulus presentations, the responses to reward were predominantly inhibitory (Wan and Peoples 2006). Yet, it is important to refer that in these electrophysiological studies no separation between D1- and D2-MSNs was performed. More recently, a key role for D2R-expressing neurons in the NAc has been identified in response to reward-predicting cue, given that these neurons increased firing at cue, but not at lever pressing, that predicted lever availability for electrical stimulation of the VTA (Owesson-White et al. 2016). Interestingly, calcium transients of D1- or D2-MSNs detected in the ventrolateral striatum with fiber photometry during a PR task, showed that D2-MSNs increase activity at trial start, but decrease as lever pressing for reward continues, reaching minimal levels at reward delivery (Natsubori et al. 2017). Moreover, higher D2-MSN calcium transients associated with cue exposure were correlated with a higher motivational state (Natsubori et al. 2017). Although dorsal and ventrolateral striatum are functionally distinct from the NAc, these data are in agreement with our results indicating that D2-MSNs in the core subregion are highly relevant for the reward-predicting cue period. Importantly, this effect may be subregion specific, because core and shell have different behavioral roles, presumably because of different inputs that they receive (Zahm 1999; Voorn et al. 2004). We focused our attention on NAc core projections to the VP because this sub-region has been proposed to be particularly relevant for acquisition of cue–reward associations, while shell seems to be more relevant for reward prediction and affective processing (Carelli 2004; Saddoris 2013; Saddoris et al. 2015; West and Carelli 2016; Sackett et al. 2017)

Another interesting finding from this study was that in reward extinction conditions, laser stimulation *per se* was not able to support the shift in preference. This data may be defying to reconcile with the fact that brief optical activation of D2-MSNs *per se* is reinforcing since it induces place preference (Cole et al. 2018). Nevertheless, these findings are similar to previous studies in which optogenetic stimulation of central amygdala (Robinson et al. 2014) or laterodorsal tegmentum-to-NAc projections (Coimbra et al. 2019) increased preference for the lever associated with optical stimulation, but only if paired with food reward. One possible explanation is that in the two-choice task, animals are mildly food restricted so their primary goal is to obtain the food pellet, so in reward extinction conditions the stimulation alone is not enough to sustain lever pressing. One other possibility is that one would need additional sessions to observe self-stimulation of these projections.

In laser extinction conditions, in which pressing either lever results in equal pellet delivery, animals pressed similarly on each lever. Still, in a reminder session where optical inhibition was associated with reward delivery in stim+ lever, animals would show a significant increase in motivation to work for that lever. Again, this suggests that D2-MSNs-VP terminals are likely suppressed during reward delivery. Although these effects could be attributed to potential alterations in the hedonic value of the reward (Berridge 2007; Berridge et al. 2009) caused by stimulation itself, the fact that stimulation during free feeding behavior did not induce any consumption differences for chow or food pellets likely rules out this hypothesis.

One possibility by which modulation of D2-MSNs in the VP could affect motivated behavior is that activation of these projections causes an indirect effect in downstream regions important for motivated behaviors, namely the VTA that is densely innervated by VP GABAergic projections (Flagel et al. 2011; Ostlund et al. 2014; Burke et al. 2017; Soares-Cunha et al. 2019). Given that optogenetic activation of D2-MSN-VP projections causes a net decrease in the firing rate of VP putative GABAergic neurons, the tonic inhibitory control that VP GABAergic neurons exert over dopaminergic neurons in the VTA is inhibited (Hjelmstad et al. 2013), which could lead to increased dopaminergic activity and consequent increase in motivated responses. Indeed, previous data from our team showed that optical activation of D2-MSNs leads to inhibition of VP GABAergic neurons, and consequently increased VTA dopaminergic activity (Soares-Cunha et al. 2019).

## Conclusion

In the present work we show that D2-MSN-VP projections differentially contribute for distinct phases of motivated behavior. Activity of these neurons is necessary to increase the value of a cue that predicts a reward, but it does not seem necessary for the operant execution of the task. This is particularly shown in the two-choice task, where, although D2-ChR2 animals show preference for one of the levers (stim-) as learning of the instrumental task progresses, the total number of lever presses (stim+ lever plus stim- lever) does not differ from control animals. Interestingly, this is also observed in studies showing preference for the stim+ lever (Robinson et al. 2014; Coimbra et al. 2019).

It is becoming increasingly evident that D2-MSNs have divergent roles in different stages of behavior, which highlights the need to perform electrophysiological recordings with opto-tagging (Kravitz et al. 2013) or calcium imaging (Klaus et al. 2017) in freely behaving animals during task performance, to determine the temporal activity of D2-MSN-VP projections during behavior.

Overall, this work shows that the role of D2-MSN-VP projections in reinforced behaviors is far more complex than anticipated, suggesting the need of a coordinated activity of the same neuronal population during behavioral performance for the execution of highly motivated behaviors.

## Materials and Methods

| Key Resources Table |  |  |  |  |
| --- | --- | --- | --- | --- |
| Reagent type (Species) or resource | Designation | Source or Reference | Identifier | Additional information |
| Strain (*Ratus norvegicus*, male) | Wistar han rat | Charles River Laboratories, Spain | NA | Wild-type rats |
| Recombinant DNA reagent | AAV5-D2R-hChR2(H134R)-EYFP | Deisseroth lab | NA |  |
| Recombinant DNA reagent | AAV5-D2R-eNpHR3.0-EYFP | Deisseroth lab | NA |  |
| Recombinant DNA reagent | AAV5-D2R- EYFP | Deisseroth lab | NA |  |
| Antibody | goat anti-GFP | Abcam | catalog #A-11055, RRID: AB_2534102 | 1:500 |
| Antibody | Alexa fluor 488 donkey anti-goat | Invitrogen | catalog #A-11055, RRID: AB_2534102 | 1:500 |
| Software, algorithm | Spike2 | Cambridge Electronic Design | RRID:SCR_000903 |  |
| Software, algorithm | Med-PC program | Med Associates | RRID:SCR_012156 |  |
| Software, algorithm | GraphPad Prism | GraphPad Software Inc | RRID:SCR_002798 | Version 8.4.0 |
| Software, algorithm | Python packages (numpy 1.10.1; scipy 0.18.1) | , Python Software Foundation, Beaverton, OR, USA | NA |  |

### Animals

Male *Wistar Han* rats (two to three months old at the beginning of the experiments) were used. Animals were maintained under standard housing conditions with 12/12h light/dark cycle (lights on from 8a.m. to 8p.m.) and room temperature of 21 ± 1°C, with relative humidity of 50–60%. Rats were individually housed after optical fiber implantation and standard diet (4RF21, Mucedola SRL) and water were given *ad libitum*, until the beginning of the behavioral experiments, in which animals switched to food restriction to maintain 85% of initial body weight.

Behavioral manipulations occurred during the light period of the light/dark cycle. Health monitoring was performed according to FELASA guidelines (Nicklas et al. 2002). All procedures were conducted in accordance with European Regulations (European Union Directive 2010/63/EU). Animal facilities and animal experimenters were certified by the National regulatory entity, Direçã o-Geral de Alimentaçã o e Veterinária (DGAV). All protocols were approved by the Ethics Committee of the Life and Health Sciences Research Institute (ICVS) and by DGAV (protocol number 19074, approved on 08/30/2016).

### Constructs and virus

eYFP or hChR2(H134R)-eYFP or eNpHR3.0-eYFP were cloned under the control of the D2R minimal promoter region as described before (Soares-Cunha et al. 2016; Zalocusky et al. 2016). Constructs were packaged in AAV5 serotype by the University of North Carolina at Chapel Hill (UNC) Gene Therapy Center Vector Core (UNC). AAV5 vector titters were 3.7–6 × 10^12^ viral molecules/ml as determined by dot blot.

### Surgery and optic fiber implantation

Rats were anesthetized with 75 mg kg^−1^ ketamine (Imalgene, Merial) plus 0.5 mg kg^−1^ medetomidine (Dorbene, Cymedica). Virus was unilaterally injected into the NAc; coordinates from bregma, according to (Paxinos and Watson 2005): +1.2 mm anteroposterior (AP), +1.2 mm mediolateral (ML), and −6.5 mm dorsoventral (DV; D2-ChR2 group and D2-eYFP control group). An optic fiber was then implanted in the VP (coordinates from bregma: −0.1 mm AP, +2.4 mm ML, and −7.5 mm DV), which was secured to the skull using two 2.4 mm screws (Bilaney) and dental cement (C&B kit, Sun Medical).

Rats were allowed to recover for three weeks before initiation of the behavioral trainings.

### Behavior

#### Animals and apparatus

Rats were habituated to 45 mg of food pellets (F0021; Bio-Serve) in the home cage, which were used as reward during the behavioral protocol, 1 day before training initiation. Behavioral sessions were performed in operant chambers (Med Associates) that contained a central, recessed magazine to provide access to 45 mg of food pellets (Bio-Serve), two retractable levers with cue lights located above them that were located on each side of the magazine. Chamber illumination was obtained through a 2.8-W, 100-mA light positioned at the top-center of the wall opposite to the magazine. The chambers were controlled by a computer equipped with the Med-PC software (Med Associates).

#### PR 1 schedule of reinforcement

All training sessions started with illumination of the house light that remained until the end of the session. On the first training session [continuous reinforcement (CRF) sessions] one lever was extended. The lever would remain extended throughout the session, and a single lever press would deliver a food pellet (maximum of 50 pellets earned within 30 minutes). In some cases, food pellets were placed on the lever to promote lever pressing. After successful completion of the CRF training, rats were trained to lever press on the opposite lever using the same training procedure. In the four following days, the side of the active lever was alternated between sessions. Then, rats were trained to lever press one time for a single food pellet in a fixed ratio (FR) schedule consisting in 50 trials in which both levers are presented, but the active lever is signaled by the illumination of the cue light above it. FR sessions began with extension of both levers (active and inactive) and illumination of the house light and the cue light over the active lever. Completion of the correct number of lever press led to a pellet delivery, retraction of the levers and the cue light turning off for a 20-s inter-trial interval (ITI). Rats were trained first with one lever active and then with the opposite lever active in separate sessions (in the same day). In a similar manner, rats were then trained using an FR4 reinforcement schedule for 4 d and a FR8 for 1 day. On the test day, rats were exposed to PR or FR experimental sessions (one session per day) according to the following schedule: day 1, FR4; day 2, PR (optical stimulation); day 3, FR4; day 4, PR (no optical stimulation). PR sessions were identical to FR4 sessions except that the operant requirement on each trial (T) was the integer (rounded down) of 1.4^(T–1)^ lever presses, starting at 1 lever press. PR sessions ended after 15 min elapsed without completion of the response requirement in a trial. All animals performed 4 sessions of PR: in the first session all animals received no optogenetic manipulation; in following sessions one third of the animals received optogenetic manipulation during ITI, one third received optogenetic manipulation at cue exposure and one third received optogenetic manipulation at pellet delivered. The days of optogenetic manipulation were counterbalanced between groups. All PR sessions were separated by two days of FR4 with no stimulation.

Optical activation consisted of: 473nm, 25ms light pulses over 1s, 10mW at the tip of the implanted fiber.

Optical inhibition consisted of: 589nm, 10 s of constant light, 10mW at the tip of the implanted fiber.

#### Two-choice schedule of reinforcement

During instrumental training, rats were presented two illuminated levers, one on either side of the magazine (Robinson et al. 2014). Presses in one lever (stim+) lead to instrumental delivery of a pellet plus laser stimulation (optical activation: 1s, 20Hz, 473nm; optical inhibition: 4s constant light, 589 nm; both with 10mW at the tip of the implanted fiber) accompanied by a 4s auditory cue (white noise or tone; always the same paired for a particular rat, but counterbalanced assignments across rats). In contrast, pressing the other lever (stim-) delivered a single pellet accompanied by another 4s auditory cue (tone or white noise), but with no laser illumination. For both levers, presses during the 4s after pellet delivery had no further consequence. Each daily session began with a single presentation of each lever (either stim+ or stim-) to ensure that the rat sampled both reward outcomes. After these two initial trials, both levers were presented simultaneously for the remainder of the session (30 minutes total), allowing the rat to freely choose between the two. Training sessions consisted in three days of Fixed Ratio (FR) 1, one day of FR4, one day of Random Ratio (RR) 4 and three days of RR6.

#### Food extinction

To assess whether laser stimulation alone could maintain responding on a stim+ associated lever when the reward was discontinued, rats were given the opportunity to earn the same levers but without pellet (pellet extinction) (Robinson et al. 2014). Each completed trial (RR4) on the stim+ lever resulted in the delivery of laser stimulation (optical activation: 1s, 20Hz, 473nm; optical inhibition: 4s constant light, 589 nm; both with 10mW at the tip of the implanted fiber) and the previously paired auditory cue but no pellet delivery. Each completed trial on the other lever (previously pellet alone) resulted in the delivery of its auditory cue but no pellet itself.

#### Laser extinction

To test the persistence of laser-induced preference, rats received 2 days reminder training with stim+ versus stim- (Robinson et al. 2014). After that, rats underwent 4 consecutive days of laser-extinction testing, where outcomes for both levers consisted in the delivery of a pellet and the associated auditory cue, with no administration of laser stimulation.

#### PR 2 schedule of reinforcement

The PR test was performed with either the stim+ lever or stim- lever (order of test conditions was balanced across animals) and was repeated for each animal with the other lever (Robinson et al. 2014). The number of presses required to produce the next reward delivery increased after each reward, according to an exponential equation (PR = [5e^(reward number*0.2)^] − 5 and rounded to the nearest integer). To determine whether any preference in responding was the result of increased workload, animals were given a FR4 session (identical to the initial day of training) after the first PR.

#### Normal chow consumption

We next examined the effect of laser stimulation on voluntary normal chow consumption in a 30-minute chow consumption test. Intake test was conducted in a familiar chamber (similar to the home cage) containing bedding on the floor in which rats had serial access to pre-weighed quantities of regular chow pellets (20 g) while also having constant access to water. Intake tests were repeated on 3 consecutive days. Laser stimulation was administered only on 1 day (optical activation: 20Hz, 473nm throughout the entire session; optical inhibition: constant light, 589 nm throughout the entire session; both with 10mW at the tip of the implanted fiber), which occurred on either day 2 or 3 (counterbalanced across rats). Control intake was measured in the absence of any laser stimulation on the 2 remaining days (day 1 and either day 2 or 3, averaged together to form a baseline measurement). Chow was weighed at the end of the test to calculate the amount consumed.

#### Food pellets consumption

We examined the effect of laser stimulation on voluntary palatable food consumption in a 30-minute test food pellet consumption test. Intake test was conducted in a familiar chamber (similar to the home cage) containing bedding on the floor in which rats had serial access to pre-weighed quantities of food pellets (around 20 g) while also having constant access to water. Intake tests were repeated on 3 consecutive days. Laser stimulation was administered only on 1 day (optical activation: 20Hz, 473nm throughout the entire session; optical inhibition: constant light, 589 nm throughout the entire session; both with 10mW at the tip of the implanted fiber), which occurred on either day 2 or 3 (counterbalanced across rats). Control intake was measured in the absence of any laser stimulation on the 2 remaining days (day 1 and either day 2 or 3, averaged together to form a baseline measurement). Food pellets were weighed at the end of the test to calculate the amount consumed.

#### Food preference

We examined the effect of laser stimulation on voluntary food preference in a 90 min free-intake test. Intake tests were conducted in a familiar chamber (similar to the home cage) containing bedding on the floor in which rats had serial access to pre-weighed quantities of chow (20 g) and food pellets (about 20 g), while also having constant access to water. Each food intake session consisted of 30 min access to 20 g of chow followed by 60 min of access to about 20 g of food pellets and chow. Intake tests were repeated on 3 consecutive days. Laser stimulation was administered only on 1 day (optical activation: 20Hz, 473nm throughout the entire session; optical inhibition: constant light, 589 nm throughout the entire session; both with 10mW at the tip of the implanted fiber), which occurred on either day 2 or 3 (counterbalanced across rats). Control intake was measured in the absence of any laser stimulation on the 2 remaining days (day 1 and either day 2 or 3, averaged together to form a baseline measurement). Chow and food pellets were weighed at the end of the test to calculate the amount consumed.

### Optogenetic manipulation

Optical manipulation was performed using either a 473 nm (ChR2) or 589 nm (NpHR) DPSS lasers, which were controlled by the MedPC software (Med Associates), through a pulse generator (Master-8; AMPI, New Ulm, MN, USA).

Before all behavioral sessions, rats were connected to an opaque optical fiber, through previously implanted ferrules placed unilaterally in the VP.

Optical stimulation was performed as follows: 473 nm; frequency of 20 Hz; 25ms pulses; 10 mW at the tip of the implanted fiber.

Optical inhibition was performed as follows: 589 nm; 4sec continuous light of 10 mW at the tip of the implanted fiber.

### *In vivo* single cell electrophysiological recordings

Three weeks post-surgery, D2-ChR2 rats (*n* = 4) were anaesthetized with urethane (1.44 g kg^−1^, Sigma). The total dose was administered in three separate intraperitoneal injections, 15 min apart. Adequate anesthesia was confirmed by the lack of withdrawal responses to hindlimb pinching. A recording electrode coupled with a fiber optic patch cable (Thorlabs) was placed in the VP (coordinates from bregma: 0 to −0.12 mm AP, +2.3 to +2.5 mm ML, and −7 to −7.6 mm DV).

Single neuron activity was recorded extracellularly with a tungsten electrode (tip impedance 5–10 Mat 1 kHz) and data sampling was performed using a CED Micro1401 interface and Spike2 software (Cambridge Electronic Design). The DPSS 473 nm laser system, controlled by a stimulator (Master-8, AMPI) was used for intracranial light delivery. Optical stimulation was performed as follows: 473 nm; frequency of 20 Hz; 12.5-ms pulses over 1 s, 10 mW.

Firing rate histograms were calculated for the baseline (10 s before stimulation), stimulation period and after stimulation period (10 s after the end of stimulation). Spike latency was determined by measuring the time between half-peak amplitude for the falling and rising edges of the unfiltered extracellular spike.

We defined the neuronal instantaneous firing rate of the *i*-th neuron as given by *r*_i_ (a_k_, b_k_) = *h* (u_i_, a_k_, b_k_, w), where *h* is a histogram function over the vector u_i_ which stores the spiking times of the *i*-th neuron in the population, within the time interval [a_k_, b_k_), and w was the bin size for *h* (usually w=1s). In order to calculate the PETH, each recorded spike train from a single neuron was aligned by the onset of optical stimulation. For each neuronal instantaneous firing rate *r*_i_ the average activity during baseline was subtracted (*r*_i_=*r*_i_-avg(*r*_i_[*t*<40s]), and then neurons were sorted by the average activity during optical stimulation.

VP GABAergic neurons were identified as those having a baseline firing rate between 0.2 and 18.7 Hz (Richard et al. 2016). Other non-identified neurons (corresponding to less that 10% of recorded cells) were excluded from the analysis.

### Immunofluorescence (IF)

Ninety minutes after initiation of the PR test, rats from Group I were deeply anesthetized with pentobarbital (Eutasil) and were transcardially perfused with 0.9% saline followed by 4% paraformaldehyde. Brains were removed and post-fixed in 4% paraformaldehyde for 24 hours. Afterwards, brains were transferred to a 30% sucrose solution (for at least 48h), and then prepared for sectioning. Coronal vibratome sections (50μm) of NAc and VP were incubated with goat anti-GFP (1:500, Abcam; catalog #ab6673, RRID: AB_305643).

Appropriate secondary fluorescent antibody was used (1:500, Invitrogen; catalog #A-11055, RRID: AB_2534102). Finally, all sections were stained with 4’,6-diamidino-2-phenylindole (DAPI; 1 mg ml^−1^).

Images were collected and analyzed by confocal microscopy (Olympus FluoViewTMFV1000). Cell counts were normalized to the area of the brain region.

### Statistical analysis

Prior to any statistical comparison between groups, normality tests (Shapiro-Wilks (S-W)) were performed for all data analyzed. When normality assumptions were met, statistical analysis using parametric tests was performed: comparison between two groups in the behavioral parameters was made using Student’s *t*-test (when normality assumptions were not met Mann-Whitney was performed instead); comparison between behavior on the ON side and the OFF side within the same subject was performed using paired t-test; Analysis of Variance (ANOVA) for repeated measures was used to compare firing rate before, during and after stimulation, and Sidak’s *post hoc* multiple comparisons was used for group differences determination (when normality assumptions were not met Friedman’s test was performed, and Dunn’s multiple comparison for post hoc analysis).

For the analysis of electrophysiological temporal variation, for each time bin, the activity during stimulation was considered significant when on that time bin the activity was out 95% of the distribution of the baseline activity. In the case of brief optogenetic stimulation datasets, given the zero mean baseline activity, the p-value was calculated as the fraction of samples in baseline activity, which were, in absolute value, greater than the value of the onset activity (single sample). For prolonged stimulation datasets, Komlogorov-Smirnov for 2 samples was performed to determine differences between the distribution of the stimulus period and the baseline.

All statistical analysis was performed using Python packages (numpy 1.10.1; scipy 0.18.1, Python Software Foundation, Beaverton, OR, USA) and GraphPad (Prism 7, La Jolla, CA, USA).

Results are presented as mean ± SEM. All of the statistical details of experiments can be found throughout the results description; these include the statistical tests used and exact p-value. The n for each experiment is indicated in the figures’ legends.

## Acknowledgments

We would like to acknowledge Karl Deisseroth from Stanford University, for providing the viral constructs.

CS-C and AJR have Scientific Employment Stimulus Contracts from Foundation for Science and Technology (FCT) (CEECIND/03887/2017; CEECIND/02130/2017). AVD has an FCT grant (SFRH/BD/147066/2019).

This work was funded by Bial Foundation 30/2016; and by FCT under the scope of the project PTDC/MED-NEU/29071/2017 (REWSTRESS).

Part of this work was developed under the scope of the project NORTE-01-0145-FEDER-000013, and FCT projects POCI-01-0145-FEDER-007038 and POCI-01-0145-FEDER-016428, supported by the Northern Portugal Regional Operational Programme (NORTE 2020), under the Portugal 2020 Partnership Agreement, through the European Regional Development Fund (FEDER) and through the Competitiveness Factors Operational Programme (COMPETE).

---

**Figure 1 – figure supplement 1.**
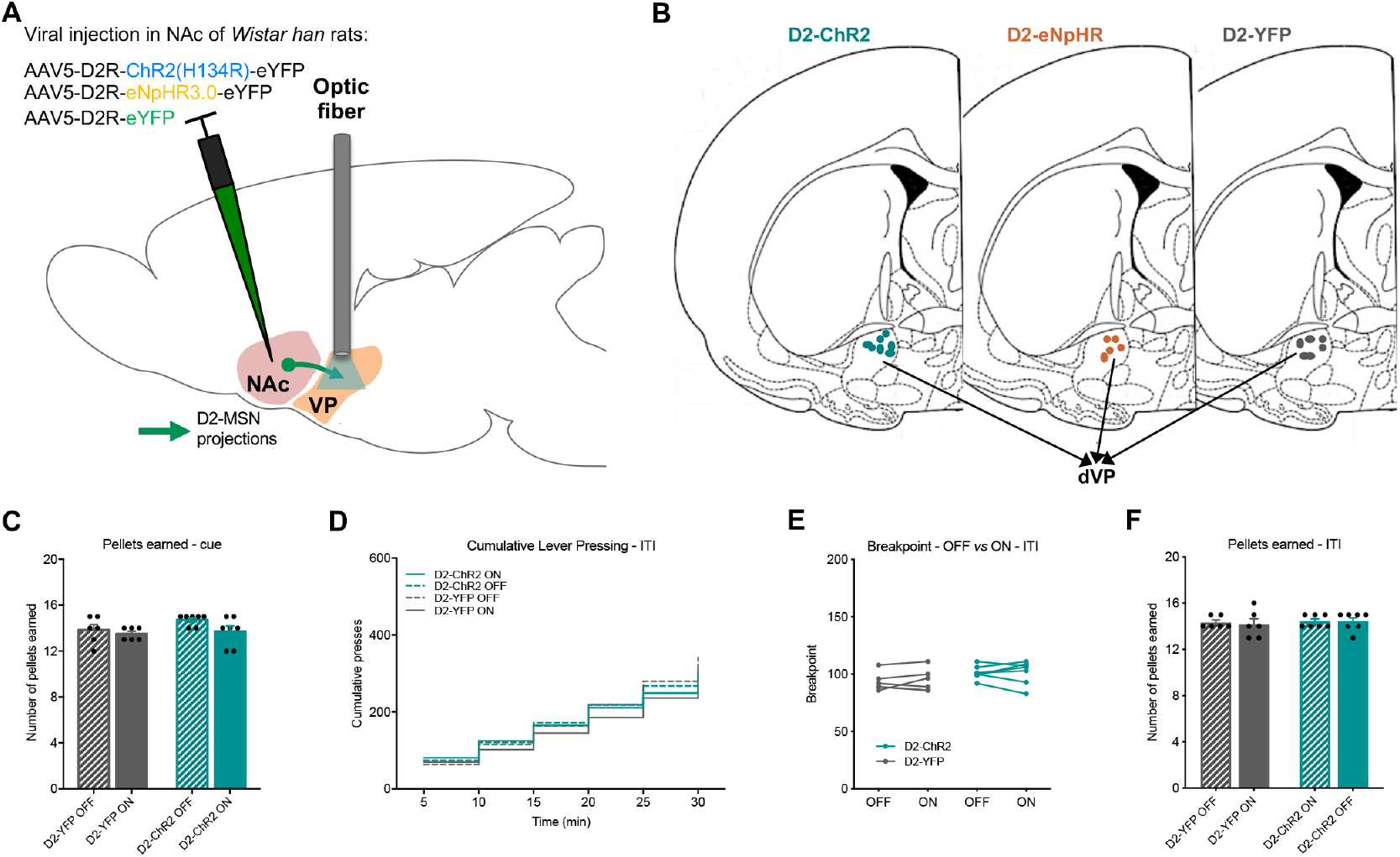
Optogenetic activation of D2-MSN-VP terminals during cue exposure increases motivation. ***A*** Strategy used for optogenetic stimulation of D2-MSN-VP terminals; rats received AAV injection of either AAV5-D2R-ChR2(H134R)-eYFP (for optical activation), AAV5-D2R-eNpHR3.0-eYFP (for optical inhibition) or AAV5-D2R-eYFP (control) in the NAc, followed by optic fiber placement in the VP. ***B*** Schematic of optic fiber placement location of D2-ChR2, D2-eNpHR and D2-eYFP rats. ***C*** In the PR session, all D2-ChR2 rats earned the same number of food pellets. ***D-F*** D2-MSN-VP terminal optical stimulation during inter-trial-interval (ITI), a period of time-out from the task, does not change ***D*** cumulative presses, ***E*** breakpoint or ***F*** the number of food pellets earned in the PR session. n_D2-ChR2_=7, n_D2-eYFP_=6. Error bars denote SEM.

**Figure 2 – figure supplement 1.**
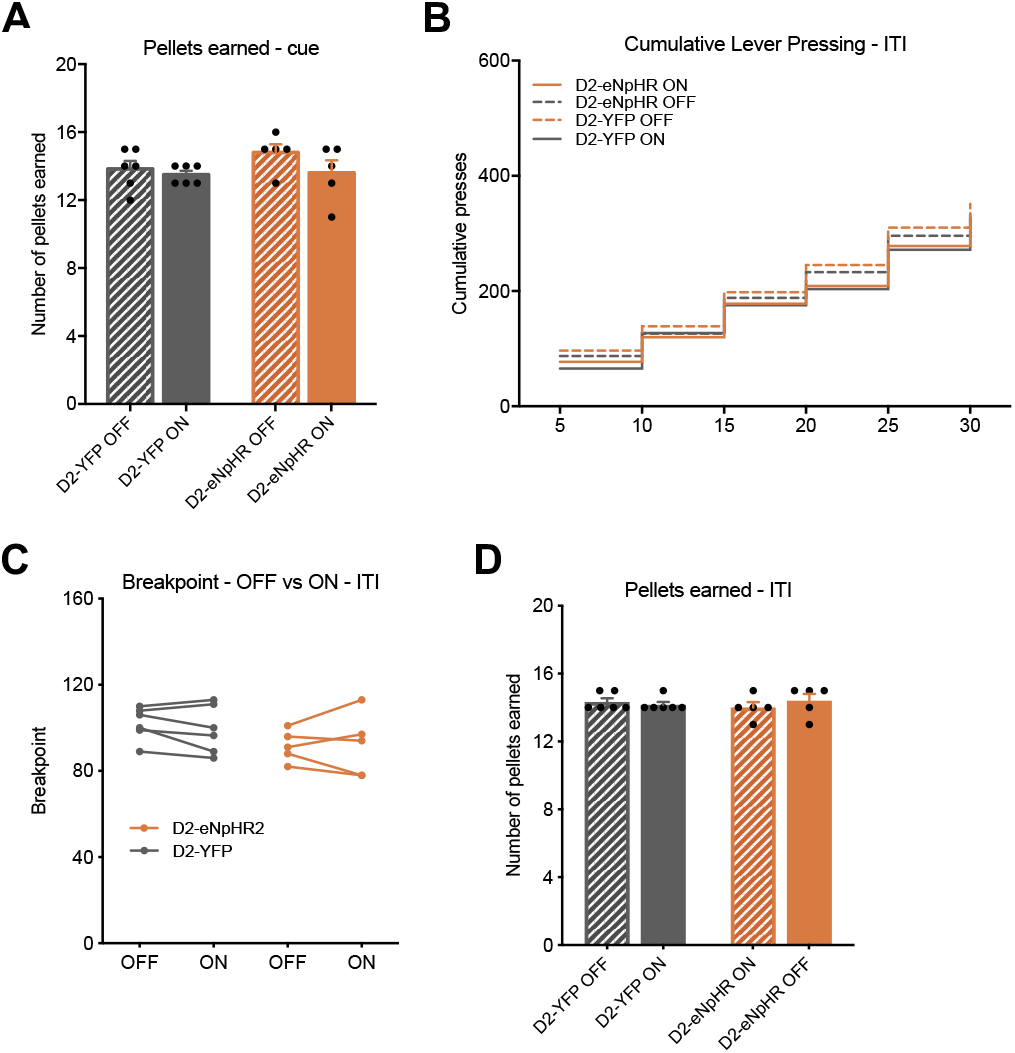
Optogenetic inhibition of D2-MSN-VP terminals during cue exposure decreases motivation. ***A*** In the PR session all D2-eNpHR rats earned the same number of food pellets. ***B-D*** D2-MSN-VP terminal optical inhibition during inter-trial-interval (ITI), a period of time-out from the task, does not change ***C*** cumulative presses, ***D*** breakpoint or ***J*** the number of food pellets earned in a PR session. n_D2-eNpHR_=5, n_D2-eYFP_=6. Error bars denote SEM.

**Figure 3 – figure supplement 1.**
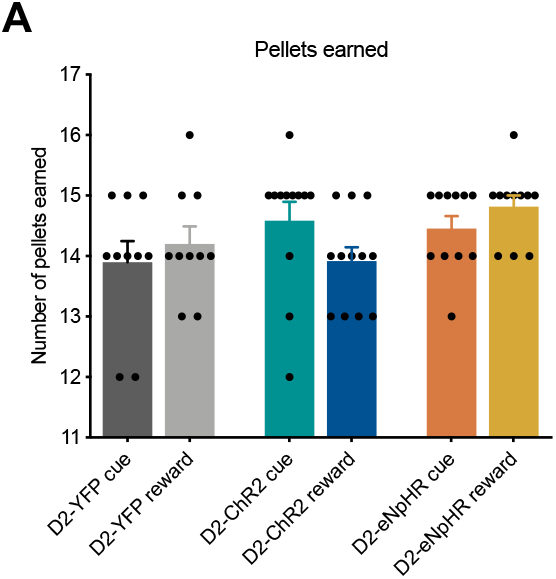
Optogenetic modulation of D2-MSN-VP terminals at reward delivery decreases motivation. ***A*** Number of food pellets earned during both PR sessions. n_D2-ChR2_=12; n_D2-eNpHR_=11, n_D2-eYFP_=10. Error bars denote SEM.

**Figure 4 - figure supplement 1.**
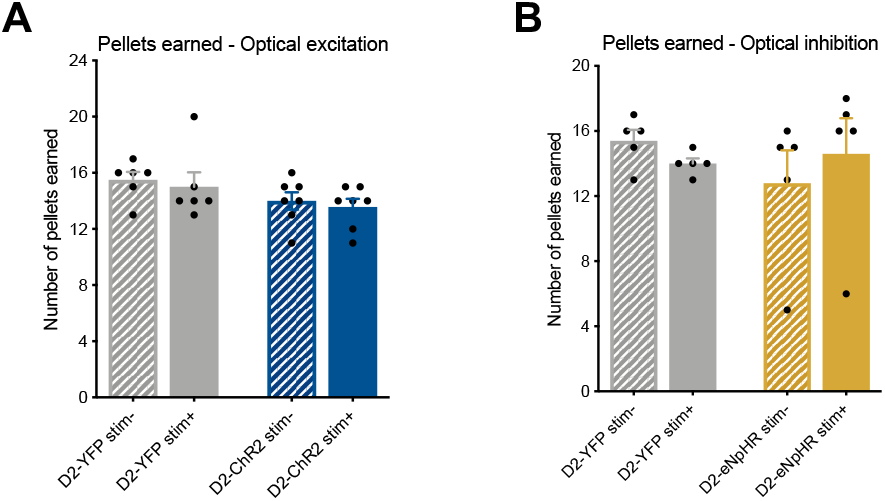
Optical activation/inhibition of D2-MSN-VP terminals paired with reward delivery reduces/increases preference and decreases motivation. ***A*** Food pellets earned during both PR sessions. ***B*** Food pellets earned during both PR sessions. n_D2-ChR2_=7, n_D2-eYFP_=6, n_D2-eNpHR_=5. Error bars denote SEM.

**Figure 4 – figure supplement 2.**
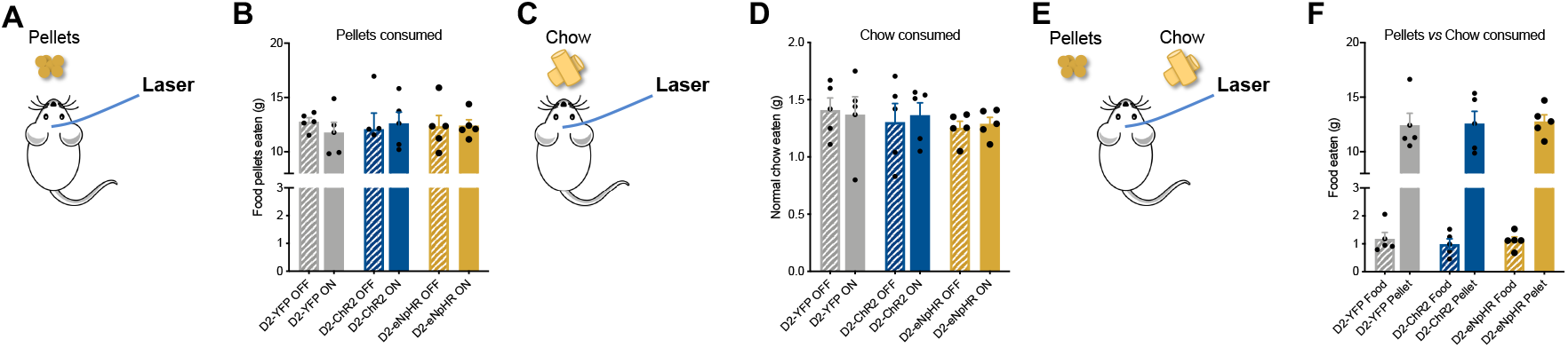
Optogenetic modulation of D2-MSN-to-VP terminals does not change food consumption. ***A*** Optogenetic modulation was performed during a free consumption behavioral session for food pellets. ***B*** The amount of food pellets consumed was similar between a session with optical modulation (ON session) and a session with no optical modulation (OFF session) for all groups. ***C*** Optogenetic activation or inhibition was given during a free consumption behavioral session for regular chow. ***D*** the amount of chow consumed was similar between a session with optical modulation (ON session) and a session with no optical modulation (OFF session) for all groups. ***E*** Optogenetic activation or inhibition was given during a free consumption behavioral session in which rats could chose to consume both food pellets and regular chow. ***F*** All rats preferred to consume food pellets, irrespective of the experimental group. n_ChR2_=5, n_D2-eNpHR_=5, n_D2-eYFP_=5. Error bars denote SEM. \*\*\**p* ≤ 0.001.

